# Role for novel family of pathogen-induced cysteine-rich transmembrane proteins in disease resistance

**DOI:** 10.1101/2020.07.25.220970

**Authors:** Marciel Pereira Mendes, Richard Hickman, Marcel C. Van Verk, Nicole Nieuwendijk, Anja Reinstädler, Ralph Panstruga, Corné M.J. Pieterse, Saskia C.M. Van Wees

## Abstract

Plants possess a sophisticated immune system to protect themselves against pathogen attack. The defense hormone salicylic acid (SA) is an important player in the plant immune gene regulatory network. Using RNA-seq time series data of *Arabidopsis thaliana* leaves treated with SA, we identified a largely uncharacterized SA-responsive gene family of eight members that are all activated in response to various pathogens or their immune elicitors and encode small proteins with cysteine-rich transmembrane domains. Based on their nucleotide similarity and chromosomal position, the designated <u>P</u>athogen-induced <u>C</u>ysteine-rich trans<u>M</u>embrane protein (PCM) genes were subdivided into three subgroups consisting of *PCM1-3* (subgroup I), *PCM4-6* (subgroup II), and *PCM7-*8 (subgroup III). Of the *PCM* genes, only *PCM4* (also known as *PCC1*) has previously been implicated in plant immunity. Transient expression assays in *Nicotiana benthamiana* indicated that most PCM proteins localize to the plasma membrane. Ectopic overexpression of the *PCMs* in Arabidopsis resulted in all eight cases in enhanced resistance against the biotrophic oomycete pathogen *Hyaloperonospora arabidopsidis* Noco2. Additionally, overexpression of *PCM* subgroup I genes conferred enhanced resistance to the hemi-biotrophic bacterial pathogen *Pseudomonas syringae* pv. *tomato* DC3000. Ectopic overexpression of the *PCMs* also affected the expression of genes related to light signaling and development, and accordingly PCM-overexpressing seedlings displayed elongated hypocotyl growth. These results point to a function of PCMs in both disease resistance and photomorphogenesis, connecting both biological processes, possibly via effects on membrane structure or activity of interacting proteins at the plasma membrane.

## INTRODUCTION

In nature and in agriculture, plants are exposed to many different pathogenic microorganisms. To counter these threats, plants have evolved a complex immune system that can perceive pathogens and activate an appropriate response. These induced defense responses aim to fortify physical barriers against pathogen entry such as callose (Luna et al., 2011). In addition, defensive compounds like secondary metabolites and pathogenesis-related proteins (PRs) accumulate, some of which have been demonstrated to possess *in vitro* antimicrobial activity and are associated with plant resistance (van Loon et al., 2006; Sels et al., 2008; Gamir et al., 2017). Plants can rely on a rich repertoire of defense compounds to combat different infecting agents. Still, many of the genes induced during pathogen infection have a so far unknown function, even though a role in defense can be expected for many of them.

The plant immune gene regulatory network that is activated in response to pathogen infection instructs which responses are expressed upon recognition of a specific invader. Conserved microbe-associated molecular patterns (MAMPs) and specific pathogen effectors can be perceived by matching receptors in the plant (Dodds and Rathjen, 2010), which subsequently activate diverse downstream signaling cascades that involve elevated levels of reactive oxygen species and calcium signaling, the modification of enzymes, and changes in hormone levels (Boller and Felix, 2009). The phytohormone salicylic acid (SA) plays a key role as signaling molecule in the regulation of plant immune responses that are primarily effective to fight biotrophic pathogens (Fu and Dong, 2013). In SA-activated cells, the transcriptional cofactor NONEXPRESSOR OF PR GENES1 (NPR1) interacts with members of the TGA family of transcription factors, leading to transcriptional activation of different other transcription factors, like members of the WRKY family, and downstream SA-responsive defense genes (Tsuda and Somssich, 2015). Microarray analysis of *Arabidopsis thaliana* (hereafter: Arabidopsis) plants expressing an NPR1-GR (glucocorticoid receptor) fusion protein (Wang et al., 2006) showed that several well-known SA-related genes, like *PR*s and *WRKY*s, were among the differentially expressed genes (DEGs) following SA treatment and dexamethasone-induced nuclear localization of NPR1. Almost 20% of the 64 direct target genes regulated by NPR1 were described as having an unknown or uncharacterized function.

While the role of SA in regulating responses to pathogen infection is well established, it is also known to have a broader influence, regulating responses to abiotic stresses, such as cold, heat shock, drought, high salinity, UV radiation, and shade avoidance (Hayat et al., 2010; Nozue et al., 2018). SA also impacts plant growth by inhibiting auxin (growth hormone) signaling and contributes to developmental processes such as flower formation. The latter is delayed in SA-deficient Arabidopsis genotypes (*NahG* transgenic lines; *eds5* and *sid2* mutants), suggesting an interplay of SA with photoperiod and autonomous (flowering) pathways (Martinez et al., 2004; Rivas-San Vicente and Plasencia, 2011).

Even though the complete Arabidopsis genome has been known for nearly two decades (Arabidopsis-Genome-Initiative, 2000), a large fraction of the protein-coding genes is still lacking a meaningful functional characterization (Niehaus et al., 2015). A common starting point for gene characterization is to reveal the conditions under which a gene is expressed. Transcriptome analysis has been extensively used to pinpoint genes that are active in specific tissue/cell types, at developmental stages or in response to different stimuli. Recently, several research groups utilized time-series transcriptome experiments in the model plant Arabidopsis to gain insight into the topology of the gene regulatory network that is engaged under different conditions. These experiments provided a wealth predictions regarding functional and regulatory roles of complete sets of genes that are differentially expressed in diverse situations (Krouk et al., 2010; Breeze et al., 2011; Bar-Joseph et al., 2012; Windram et al., 2012; Lewis et al., 2015; Coolen et al., 2016; Hickman et al., 2017). In our recent study, we applied whole transcriptome shotgun sequencing (RNA-seq) time series and found that approximately one-third of the Arabidopsis genome was differentially expressed in leaves upon treatment with SA over a 16-h time course, with changes in gene expression occurring in well-defined process-specific waves of induction or repression (Hickman et al., 2019).

Here, this SA-responsive gene set was analyzed with the comparative genomics tools OrthoMCL and JackHMMER, which identified homologous groups of largely uncharacterized genes that may play a role in SA-associated immunity. This integrated analysis categorized over a hundred groups of SA-responsive genes, including one group of eight genes encoding short proteins that share a predicted cysteine-rich transmembrane domain and are also responsive to various pathogens and immune elicitors. We therefore named them <u>P</u>athogen-induced <u>C</u>ysteine-rich trans<u>M</u>embrane proteins (PCMs). The *PCM*s are also present in the group of NPR1-regulated direct target genes, mentioned above. Cysteine-rich repeat proteins have been predicted to be involved in biotic and abiotic stress responses (Venancio and Aravind, 2010). For one of the family members (PCC1/PCM4) a role as positive regulator of defense to the biotrophic pathogen *Hyaloperonospora arabidopsidis* has been demonstrated (Sauerbrunn and Schlaich, 2004), while for another family member (CYSTM3/PCM8) a role as negative regulator of salt stress responses has been reported (Xu et al., 2019). Analysis of Arabidopsis PCM-overexpressing lines revealed a positive role of these proteins in immunity against pathogens with (hemi)biotrophic lifestyles. Furthermore, we expanded the potential scope of their function to a role in photomorphogenesis and hypocotyl development.

## RESULTS

### Analysis of uncharacterized SA-responsive genes identifies a family of cysteine-rich transmembrane proteins

Recently, we used high-throughput RNA-seq analysis to profile genome-wide changes in mRNA abundance in Arabidopsis leaves following treatment with SA over a 16-h period. Analysis of these transcriptome data identified 9524 genes that were differentially expressed between mock-and SA-treated leaves (Hickman et al., 2019). Subsequent investigation of functional annotations associated with these differentially expressed genes (DEGs) revealed that 630 of these genes encode proteins of unknown or uncharacterized function. Because of the central role of SA in defense against pathogen infection we hypothesized that among these genes would be genes with undiscovered roles in plant immunity. To simplify the analysis and functional interpretation of these uncharacterized genes, we first divided them in groups based on amino acid sequence similarity. To achieve this, we used OrthoMCL (Li et al., 2003), which is a tool for identifying homologous relationships between sets of proteins. This analysis resulted in a division of 101 groups of putative homologs, each comprising between two and nine members (Supplemental Data set 1; Figure 1A). Because we were specifically interested in genes that are involved in defense against pathogens, we analyzed gene behavior, using available gene expression data from Genevestigator (http://www.genevestigator.ethz.ch/) (Hruz et al., 2008). This pointed to a group of seven genes that were highly induced by a variety of immune elicitors and pathogens (Figure 2A) and that were responsive to SA in our RNA-seq experiment (Figure 2B).

**Figure 1.**
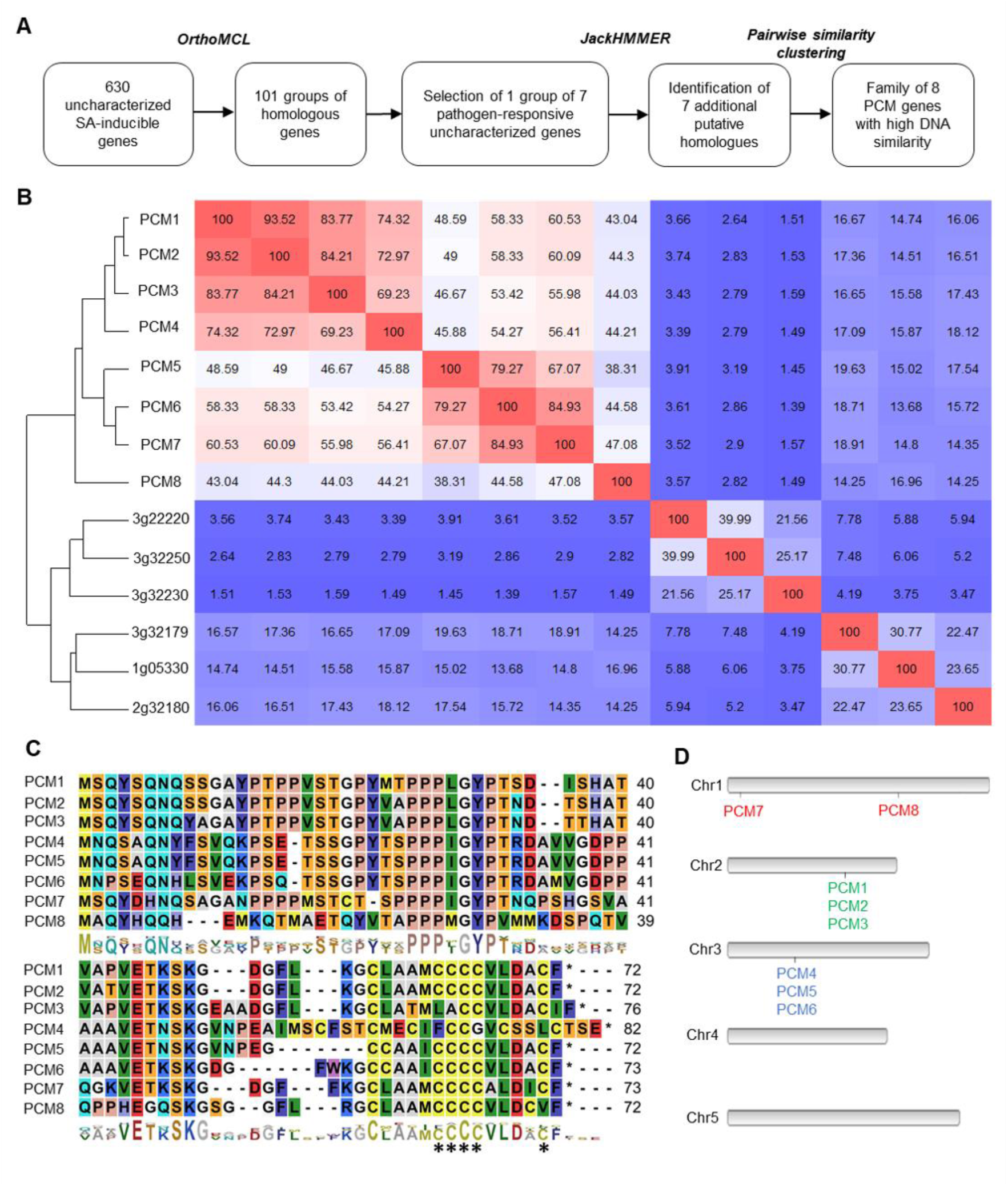
Identification of groups of homologous, uncharacterized SA-inducible genes; Selection of the *PCM* gene family. (**A**) Workflow to identify groups of homologous, unknown SA-inducible genes. First, SA-induced DEGs were grouped by DNA similarity using OrthoMCL. One pathogen-responsive group was subjected for further analysis using JackHMMER, followed by pair-wise similarity clustering, revealing a distinct family of eight homologous *PCM* genes. (**B**) DNA similarity matrix showing the 14 genes identified by the JackHMMER search. Red and blue indicate high and low similarity, respectively. Unsupervised hierarchical clustering identified a distinct group of *PCM* genes with high DNA similarity. (**C**) Amino acid sequence alignment of the eight PCMs. The conserved cysteine-rich transmembrane domain (CYSTM) is highlighted. (**D**) The locations of the eight *PCM* genes on the Arabidopsis chromosomes (Chr1 to Chr5). The different color gene names reflect *PCM* distribution across chromosomes.

**Figure 2.**
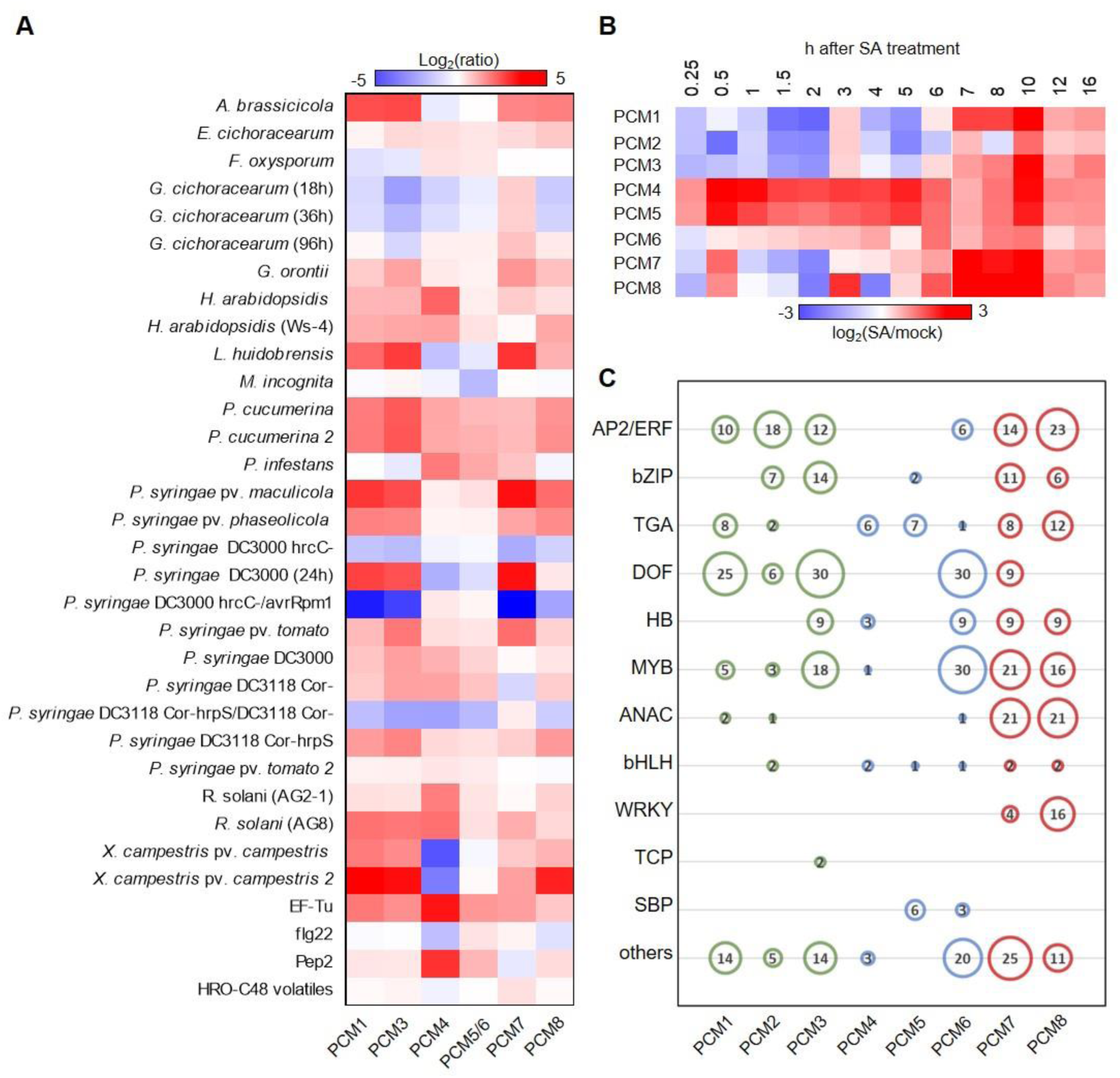
Expression behaviour of *PCM* genes. (**A**) Genevestigator expression analysis. Shown is a heatmap of expression ratios for the *PCM* genes following treatments with biotic stressors/elicitors. On the microarrays from which these data are derived (*P* < 0.001), probes for *PCM2* are missing, and the probes for *PCM5* and *PCM6* are shared. (**B**) Temporal expression of *PCM* genes over a 16-h time course upon exogenous application of SA. Red and blue indicate increased and decreased expression, respectively. (**C**) Representation of DNA-binding motifs in the promoters of the *PCM* genes. Motifs are grouped according to cognate transcription factor family. The size and number in each circle represent the per-family motif count.

To identify all possible paralogs (including remote paralogs), the seven genes were used as queries in JackHMMER (Finn et al., 2015) (www.ebi.ac.uk/Tools/hmmer/search/jackhmmer). JackHMMER is a highly sensitive homology detection tool that can identify shared protein domains among matched sequences, as defined according to Pfam domains (Finn et al., 2015). This analysis led to the prediction of seven additional paralogs (Supplemental Data set 2). Next, we quantified the degree of nucleotide sequence identity between the 14 proteins by constructing a nucleotide sequence identity matrix (Figure 1B), which was followed by unsupervised clustering of the similarity matrix, leading to the identification of a distinct family of eight small genes (<82 amino acids (AA)) with high nucleotide sequence identity (>38%). All of the seven originally selected genes of unassigned function belong to this group, including *PCC1* (*PCM4* in Figure 1), which has a reported role in defense and is regulated by the circadian clock (Sauerbrunn and Schlaich, 2004). One other member, *CYSTM3* (*PCM8* in Figure 1), has very recently also been characterized and shown to negatively influence salt stress resistance (Xu et al., 2019). Supplemental Table 1 lists all the *PCM* genes with their AGI number and alternative name. The genes in this family all encode short proteins (71-82 AA) with a conserved cysteine-rich transmembrane (CYSTM) domain, as predicted by the JackHMMER analyses (Figure 1C). To reflect their regulation and enrichment for cysteine residues in the encoded proteins, this eight-member gene family was named pathogen-induced cysteine-rich transmembrane proteins (PCMs). The *PCM* gene family contains two distinct gene clusters; the *PCM1, PCM2* and *PCM3* genes are situated in tandem on Arabidopsis chromosome 2, while *PCM4* (*PCC1*), *PCM5* and *PCM6* are tandemly arrayed on chromosome 3 (Figure 1D). Furthermore, *PCM7* and *PCM8* (*CYSTM3*) are positioned at distant locations on chromosome 1. The expression behavior of the eight *PCM* genes is broadly along the lines of the three subgroups, showing overlap but also differences with members of the other subgroups (Figure 2A and 2B). This is in accordance with varying overrepresentation of different transcription factor-binding DNA motifs in the promoters of the eight *PCM* genes (Figure 2C). The remainder of this paper explores the significance of the PCM protein family and its three subgroups in plant immunity and development.

### Subcellular localization of PCMs

The characteristic CYSTM domain that resides in the PCM protein family is encoded by a total of 98 genes across 33 plant species (Supplemental Figure 1). Transmembrane domains enable protein functions across membranes (Luschnig and Vert, 2014) (Sharpe et al., 2010) and are often conserved across kingdoms when the respective protein has a specialized function (e.g., photoreceptors in eyes of mammals and insects) (Fischer et al., 2004). To begin to characterize the PCMs, we determined their subcellular localization by fusing the Venus yellow fluorescent protein (YFP) to the C-terminus of all eight PCM proteins and expressing these fusion proteins under the control of the constitutively active cauliflower mosaic virus (CaMV) 35S promoter. Expression of empty vector (EV-YFP, resulting in free YFP) served as a control, and the dye FM 4-64 was used as a membrane marker. Confocal microscopy analysis of *Agrobacterium*-infiltrated *Nicotiana benthamiana* leaves transiently expressing the fusion proteins, confirmed plasma membrane localization for five of the PCM-YFP variants (Figure 3). In case of the PCM1, PCM2, PCM3, PCM4 and PCM5 fusion proteins, the YFP signal overlapped with the fluorescent signal of the plasma membrane-localized FM 4-64 dye, suggesting plasma membrane localization of these proteins. By contrast, in case of the YFP-tagged family members PCM6, PCM7 and PCM8 the YFP signals were detected prevalently in the cytoplasm and the nucleus, which could either be indicative of a non-membrane localization of these proteins or reflect undesired cleavage of the YFP label.

**Figure 3.**
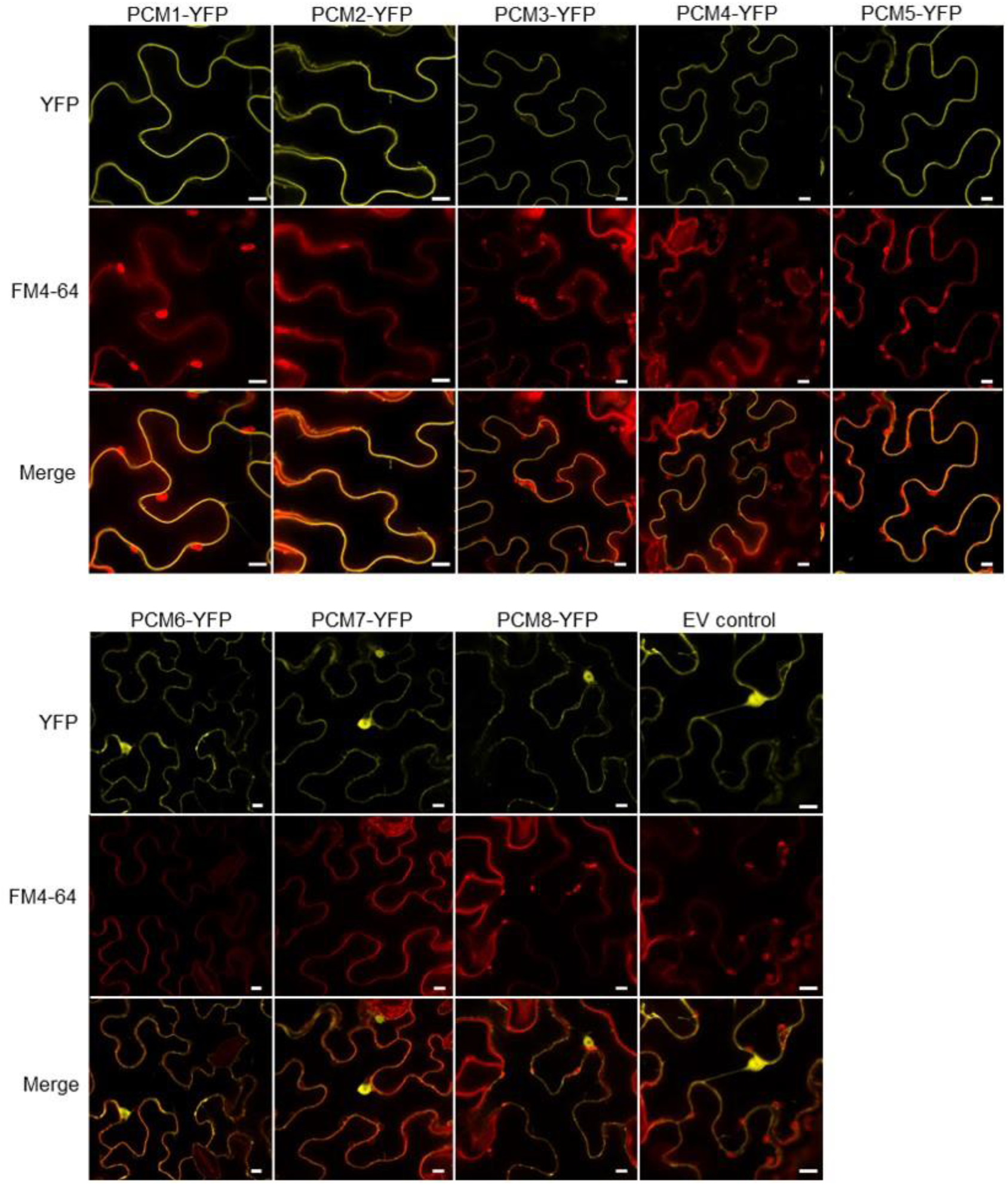
Subcellular localization of PCM-YFP fusion proteins. Confocal images of transiently transformed *N*. *benthamiana* epidermal leaf cells expressing the eight YFP-tagged PCM proteins under control of the CaMV 35S promoter. Representative fluorescence images are shown of PCM-YFP or free YFP (control) in the top panels, of FM 4-64 labelling of the membranes in the middle panels, and of the overlay of YFP and FM 4-64 in the bottom panels. Bar = 10 μm.

### *PCM* coexpression analysis points to specificity in PCM function

Because genes with related biological functions often have similar expression patterns, a well-established method to investigate gene function is the construction and analysis of gene coexpression networks (Vandepoele et al., 2009). Using the eight *PCM* genes as query we generated *PCM* coexpression networks using publicly available microarray and RNA-seq datasets with the ATTED-II coexpression tool (Obayashi et al., 2017) (Figure 4). The *PCM* coexpression network was enriched for genes associated with defense responses (*P <* 0.01; hypergeometric test) and included known defense-related genes such as *LURP1, ACD6, RLP36, NTL6, NAC61, NAC90, ZFAR, PDR12, WRKY75* and *MPK11*, suggesting a role for the PCM protein family in plant defense. Within the *PCM* coexpression network, coexpression neighborhoods of members of the three *PCM* sub-groups (Figure 2) overlap. Interestingly, the coexpression neighborhood occupied by subgroup II (*PCM4, PCM5* and *PCM6*) was distinct from that of all other *PCM* genes. Also, *PCM7* was part of a relatively isolated coexpression subnetwork. On the other hand, *PCM8* shared its coexpression neighborhood to a large extent with that of subgroup I (*PCM1, PCM2* and *PCM3*). In sum, our coexpression network analysis suggests a role for all eight *PCM* genes in plant defense, but also highlights subnetworks, suggesting functional diversification and/or differential regulation of the *PCM* subgroups. This notion is further supported by the distinct gene expression behavior of the different *PCM* subgroups after treatment with pathogens or exogenous SA and the presence of different transcription factor binding sites in the promoters of the *PCM* genes (Figure 2C).

**Figure 4.**
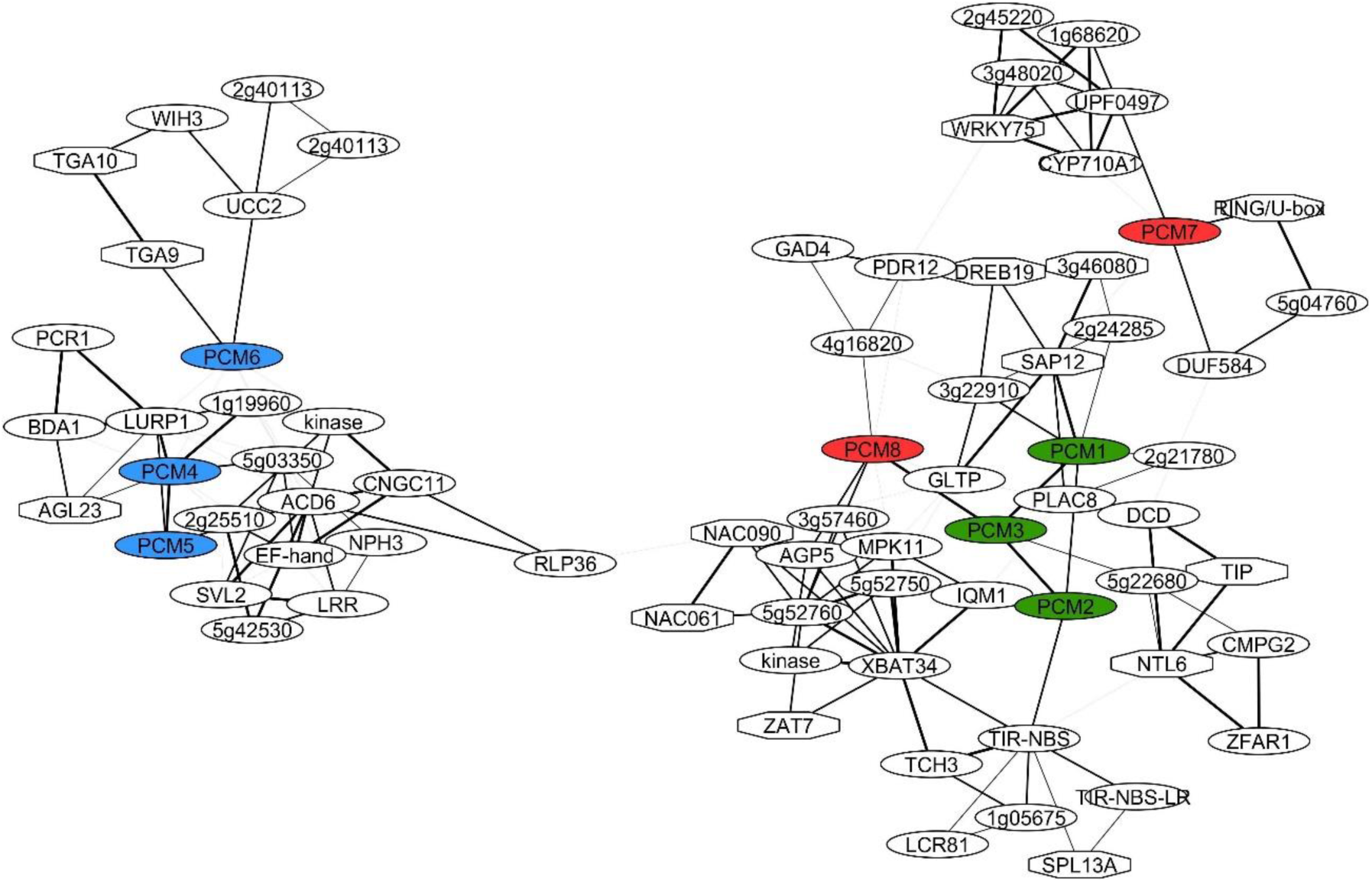
*PCM* coexpression networks. Coexpression network obtained using the ATTED-II Network Drawer tool on whole-genome transcriptome data sets with *PCM* genes as bait. Hexagonal-shaped nodes indicate genes encoding transcriptional regulators. The thickness of the lines is proportional to the extent of coexpression of the linked gene.

### PCM-overexpressing lines show enhanced resistance to (hemi)-biotrophic pathogens

To investigate the hypothesis that members of the PCM protein family play a role in plant immunity, transgenic Arabidopsis lines expressing the individual *PCM* genes under the control of the CaMV 35S promoter were generated. The transgenic *PCM*-overexpression (*PCM*-OX) lines were of unaltered size and did not show any obvious developmental abnormalities (Supplemental Figure S2). RNA-seq analysis (see below) confirmed the overexpression status of the *PCM*-OX lines for genes *PCM1* and *PCM7*, but not for *PCM5* whose overexpression levels might have remained below the thresholds of statistical analysis (Supplemental Data set 2). Because the *PCM* gene family responded to exogenous SA treatment (Figure 2B), the *PCM*-OX lines were screened for an altered level of resistance to two pathogens that are controlled by SA-dependent defenses: the obligate biotrophic oomycete *H*. *arabidopsidis* Noco2 (*Hpa* Noco2) and the hemi-biotrophic bacterium *P*. *syringae* pv. *tomato* DC3000 (*Pto* DC3000). For both assays, the performance of 5-week-old *PCM*-OX lines was compared to that of the wild-type (Col-0) and the enhanced susceptible mutant *eds1*-2 of the same age. With the exception of *PCM6*, overexpression of all other *PCM* genes led to reduced *Hpa* Noco2 spore formation when compared to wild-type plants (Figure 5A). *Pto* DC3000 propagation was significantly decreased in the *PCM1*-OX, *PCM2-*OX, and *PCM3-*OX lines but not in the other lines (Figure 5B). These findings suggest that the vast majority of PCM family members is positively involved in host defense against *Hpa* Noco2, while a protective effect against *Pto* DC3000 is only evident for the subgroup I of the PCM protein family comprising PCM1, PCM2, and PCM3.

**Figure 5.**
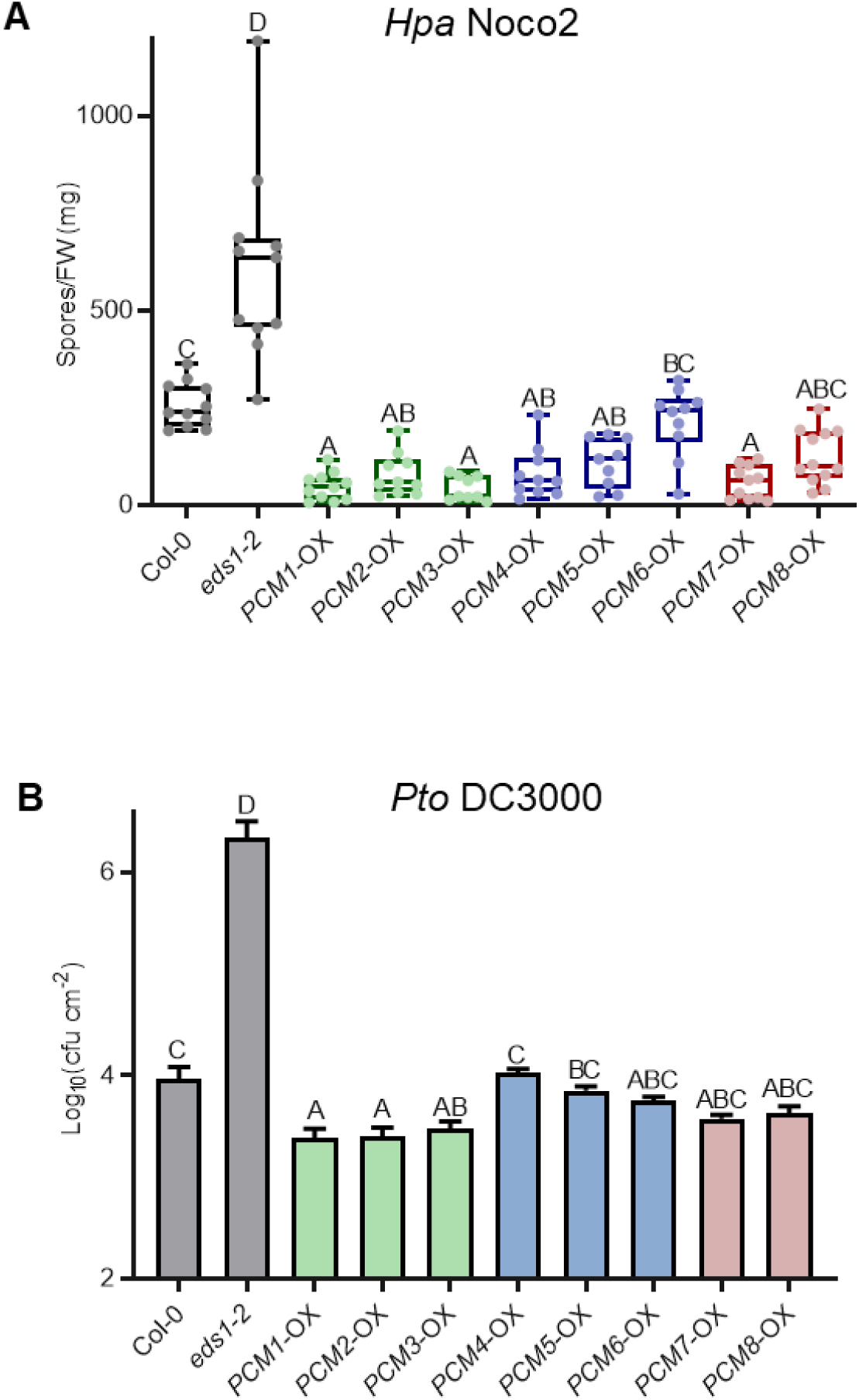
Overexpression of *PCMs* enhances resistance to *Hpa* and *Pto* DC3000. (**A**), Quantification of *Hpa* Noco2 sporulation on 5-week-old wild-type (Col-0), *eds1*-2 and transgenic lines constitutively overexpressing individual *PCM* genes under the control of the CaMV 35S promoter (*PCM*-OX) at 10 days post inoculation (dpi) by spraying (*n* = 9-12). (**B**), Bacterial multiplication of *Pto* DC3000 in wild-type (Col-0), *eds1*-2 and *PCM*-OX lines at 3 dpi by pressure infiltration (*n* = 8). Means ± SE (error bars) are shown. Letters denote significant differences between genotypes (one-way ANOVA, Tukey’s post-hoc test, *P* < 0.05). Experiment repeated with similar results.

### Transcriptome analysis of *PCM*-OX lines reveals no upregulation of typical immune responses

To gain insight into the mechanisms underlying the enhanced disease resistance phenotype obtained by overexpression of the *PCM* genes, we analyzed the transcriptome of three *PCM*-OX lines, each representing a member of the three PCM subgroups: *PCM1*-OX (subgroup I), *PCM5*-OX (subgroup II), and *PCM7*-OX (subgroup III). RNA-seq analysis was performed on leaf tissue harvested from 5-week-old, non-treated plants. Differential expression analysis revealed that in the *PCM1-*OX, *PCM5-*OX and *PCM7-*OX lines 934, 873, and 515 genes, respectively, were differentially expressed in comparison to wild-type Col-0 plants (*P* < 0.05, fold change > 2) (Supplemental Data set 2). Among the DEGs there were *PCM1* in the *PCM1*-OX line and *PCM7* in the *PCM7*-OX line, each showing a 2-fold log increase in transcript abundance. Notably, the list of DEGs comprised no other *PCM* gene in any of the overexpression lines, indicating that there is no compensatory regulation of other family members in this situation. There was considerable overlap between the expression profiles of the three *PCM-*OX lines (Figure 6A). Of all DEGs, 27% were upregulated or downregulated in all three lines (in the same direction), whereas 44% were specifically up-or downregulated in a single overexpression line (Figure 6B). More genes were downregulated (60%) than upregulated (40%).

**Figure 6.**
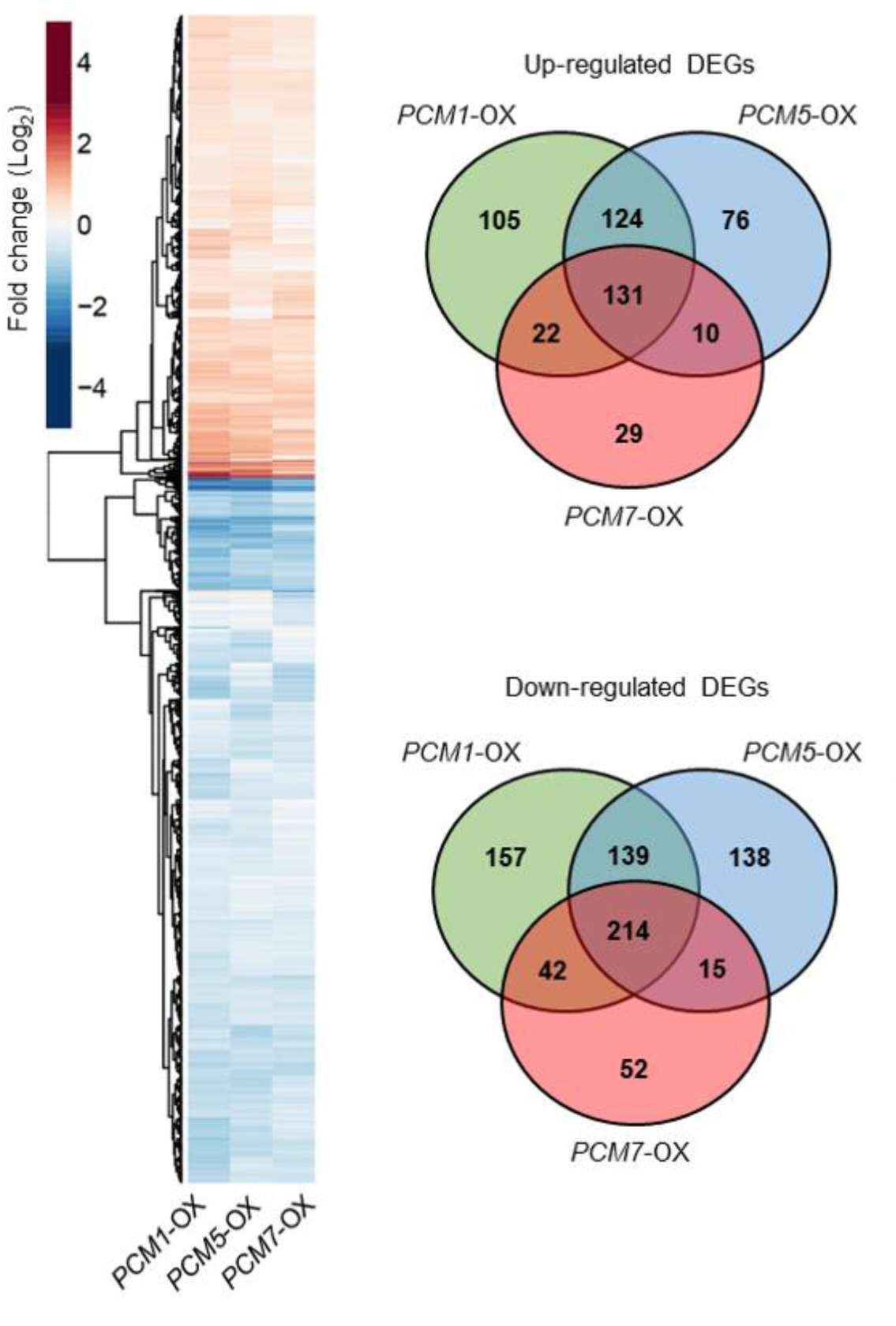
Transcriptome analysis of *PCM*-OX lines. (**A**), Heatmap (left) showing up-and downregulation of genes in the *PCM1*-OX, *PCM5*-OX or *PCM7*-OX lines in comparison to wild-type Col-0 plants, as revealed by RNA-seq analysis. (**B**), Venn diagrams (right) indicating the overlap between DEGs in each of the *PCM*-OX lines.

The overlapping 131 upregulated DEGs shared by all three *PCM-*OX lines were not enriched for typical immunity-related functions (Figure 7). Instead, the term ‘circadian rhythm’ was the most significantly enriched specific category, with additional enriched terms including ‘regulation of multicellular organismal development’ ‘plant cell wall loosening’, and ‘response to red or far red light’. The shared 214 downregulated DEGs by all three *PCM-*OX lines were associated with functional categories such as ‘rRNA processing’, ‘response to cytokinin’ and ‘response to light stimulus’ (Figure 7). There was also no enrichment of purely immunity-related categories DEGs that were specifically up-or downregulated in any of the *PCM1-*OX, *PCM5-*OX or *PCM7-*OX lines. General terms like ‘response to hormone’ were overrepresented in different lines though, while ‘glycosinolate process’ was specifically enriched in up-regulated DEGs of *PCM7*-OX, and ‘nucleolus’ was overrepresent in *PCM7*-OX (up- and downregulated) and *PCM5*-OX (only downregulated). Based on these data, we hypothesize that pathogen-induced PCM production contributes to an increased level of defense through an impact on developmental processes in the cells that may affect pathogen performance.

**Figure 7.**
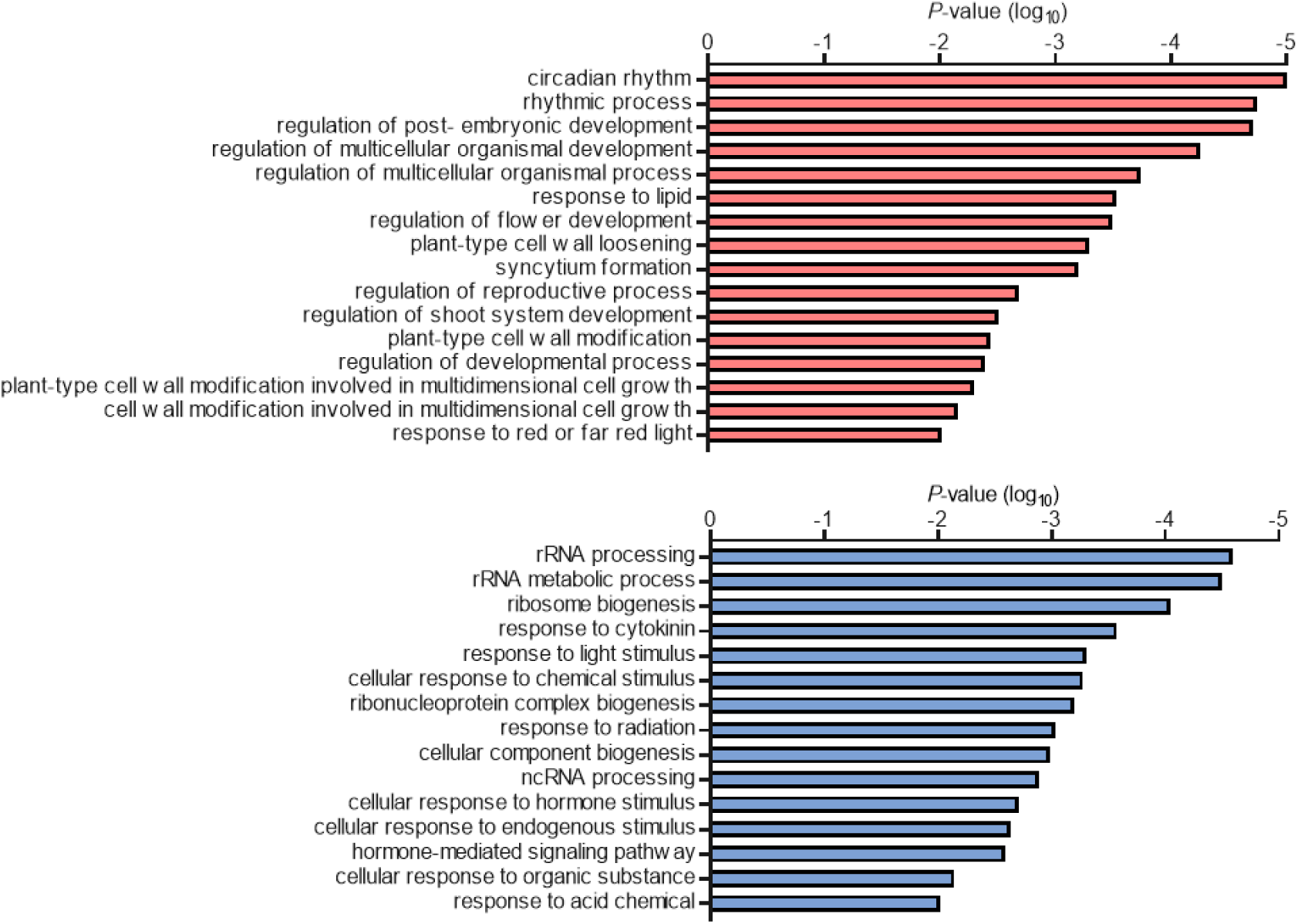
GO terms enriched among genes up- or downregulated in *PCM*-OX lines. Shown are the GO terms significantly enriched among the genes that are significantly upregulated (top) or downregulated (bottom) in *PCM1*-OX, *PCM5*-OX and *PCM7*-OX, when compared to wild type.

### Involvement of PCMs in hypocotyl elongation

There was no clear link to plant immunity among the genes differentially expressed in the three *PCM*-OX lines assayed, while the association with developmental processes and light responses was obvious (Figure 7). In all three *PCM-*OX lines, the *HY5* and *HYH* genes, which are master regulators of light signaling and also respond to pathogen infection (Genevestigator data), were upregulated. This prompted us to investigate morphogenic responsiveness of the *PCM*-OX lines. In shade-avoiding plants such as Arabidopsis, perception of far-red light triggers morphological adaptations such as elongation of the hypocotyl and petioles in order to reach for better quality light (Ballaré, 2014). The *hy5 hyh* double mutant, which is affected in HY5 and its closely related HY5 homolog (HYH), displays such elongated hypocotyl growth compared to wild-type plants when cultivated in white light (Van Gelderen et al., 2018). Unexpectedly, the hypocotyl length of the *PCM1-*OX, *PCM5-*OX, and *PCM7*-OX lines was also greater than that of the wild type, and the hypocotyl of *PCM7*-OX was even of the same size as that of the *hy5 hyh* double mutant (Figure 8). This points to a role for PCMs in modulating both growth and development. Possibly, the PCMs affect HY5 protein activity or stability, which is compensated by an enhanced expression level of the *HY5* gene. Altogether, our data suggest dual roles for PCMs in defense and in photomorphogenesis.

**Figure 8.**
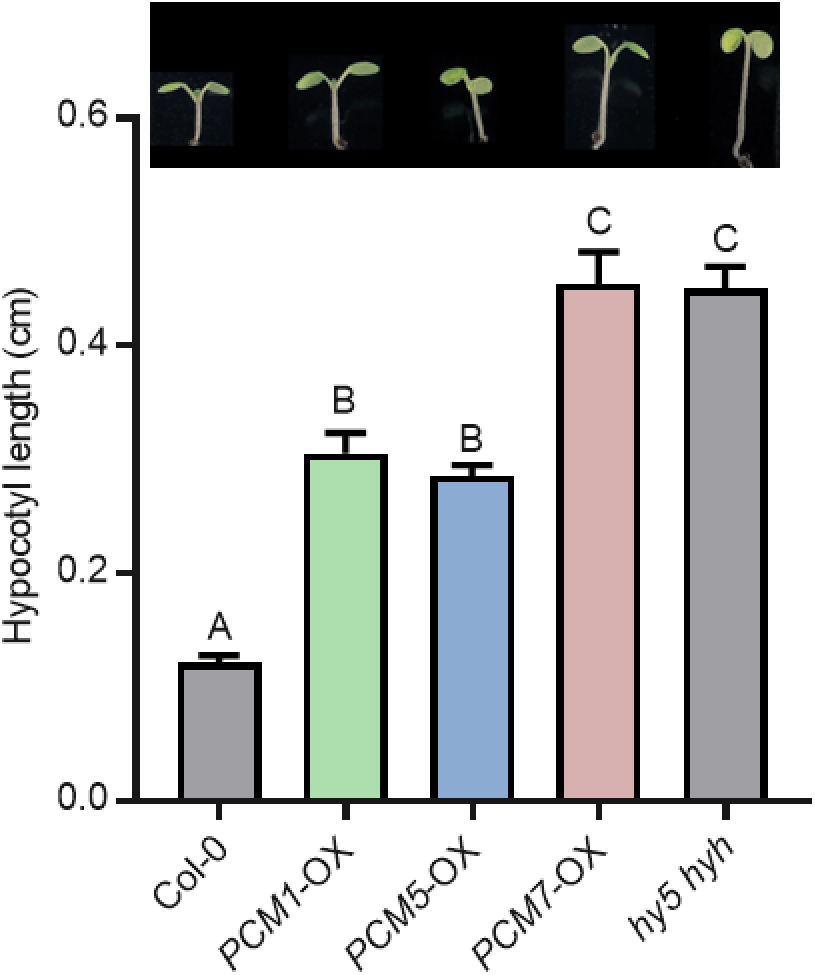
*PCM1, PCM5*, and *PCM7* influence hypocotyl elongation. Hypocotyl lengths of 7-day-old Col-0, *PCM1*-OX, *PCM5*-OX, *PCM7*-OX and *hy5 hyh* seedlings grown *in vitro* in white light (*n* = 20). Means ± SE (error bars) are shown. Letters denote significant differences between genotypes (one-way ANOVA, Tukey’s post-hoc test, *P* < 0.05). Inset: representative pictures of 7-day-old seedlings. Experiment repeated with similar results.

## DISCUSSION

Despite over two decades of research efforts focused on the model plant Arabidopsis, a significant fraction (over 13%) of genes found in this plant are not characterized to any extent (Luhua et al., 2013; Niehaus et al., 2015). Our analysis of SA-responsive genes in Arabidopsis leaves revealed that 630 genes encode proteins of unknown function. Using a protein homology search we grouped these uncharacterized genes into 101 groups of paralogs that likely encode proteins with similar functions (Figure 1A; Supplemental Data set 1). We validated whether such an approach could aid the functional annotation of groups of unknown genes. Therefore, we selected and further characterized a family of eight pathogen-induced cysteine-rich transmembrane proteins (PCMs). The *PCM* genes formed three subgroups, based on their nucleotide similarity and chromosomal position (Figure 1B and 1D). The expression profiles of the *PCM* members under different biotic stress conditions and SA treatment broadly followed that of the three subgroups, showing some overlap but also differences between the subgroups (Figure 2A and 2B). This is in agreement with the commonalities and dissimilarities in transcription factor binding sites detected in the promotor regions of the eight *PCM* genes (Figure 2C), and the overlap or isolation of the coexpression networks of the *PCM* genes (Figure 4).

Given the complexity of the *PCM* family and its overlap in coexpressed genes, we expected functional redundancy between members and thus resorted to using overexpression lines rather than knockout mutants for the functional analysis of this protein family. Overexpression of one *PCM* member of each of the three subgroups (*PCM1*-OX, *PCM5*-OX and *PCM7*-OX) revealed 27% overlap of all DEGs, and 44% specific expression by one of the *PCM* genes (Figure 6). A function for the PCMs in defense was evidenced by the enhanced resistance of *PCM* overexpression lines to the biotrophic pathogens *Hpa* Noco2 and *Pto* DC3000 (Figure 5). Moreover, *PCM* overexpression resulted in differential expression of genes related to light and development, and seedlings displayed elongated hypocotyl growth, suggesting an additional role for PCMs in photomorphogenesis (Figure 8). Though we only used single *PCM*-OX lines in our sets of experiments, the shared phenotypes regarding DEGs, pathogen resistance and photomorphogenesis suggest that these effects are authentic consequences of *PCM* overexpression and not the result of position effects of the transgenes.

### Membrane association of CYSTM domain-containing PCMs

The PCMs are small proteins (<84 AA) that contain a predicted cysteine-rich transmembrane C-terminus domain (CYSTM), which is a rare domain, but highly conserved among eukaryotic organisms. CYSTM domain-containing proteins are present in diverse species, including Arabidopsis, *Caenorhabditis elegans, Candia albicans, Homo sapiens, Mus musculus, Oryza sativa, Saccharomysces cerevisae* and *Zea mays* (Venancio and Aravind, 2010). The molecular mechanism by which the CYSTM module functions is not clear yet, but the proteins appear to play a role in stress tolerance, for example by altering the redox potential of membranes, thereby quenching radical species to protect the plant, or by affecting membrane-associated protein functions (Kuramata et al., 2009; Venancio and Aravind, 2010).

The conserved cysteines may serve to interact with a ligand, e.g. other PCMs, which could result in homo- or heterodimerization as shown in yeast expression systems for several Arabidopsis PCM family members (Mir and Leon, 2014; Xu et al., 2018). However, the PCMs can potentially also interact with other protein partners, as shown for PCM4/PCC1, which interacts with its N-terminal part (cytoplasm-faced, non-CYSTM containing) with the subunit 5 of the COP9 signalosome at the plasma membrane. This may lead to post-translational control of multiple protein targets involved in diverse biological processes such as light signaling, development, and immunity (Mir et al., 2013; Mir and Leon, 2014).

We experimentally confirmed a tight association of PCM1-PCM5 with the cell periphery and the fluorescent FM4-64 marker (Figure 3), suggesting that these proteins are anchored to the plasma membrane. This localization could potentially promote a change in local lipid composition, as shown for PCM4/PCC1 (Mir et al., 2013), and also affect the membrane structure. This notion is supported by changes in gene expression observed in the *PCM*-OX lines, which highlighted enrichment in gene ontology (GO) terms related to ‘response to lipid’, cell wall modification’, and ‘regulation of development’ (Figure 7). Moreover, membrane alterations may block the invasion of intracellular pathogens like *Hpa* (Figure 5A) that form an intricate interface with the host membrane. There may also be consequences for membrane permeability or activity of (defense regulatory) proteins associated with the plasma membrane. While plasma membrane localization of PCM1-PCM5 was supported by experiments using YFP-tagged PCMs in transiently transformed *N*. *benthamiana* leaves, this was not the case for PCM6-PCM8 (Figure 3). The latter finding is consistent with a recent study by Xu et al. (2018) who also found cytoplasmic localization of these proteins using the same study system. However, these authors also reported cytoplasmic localization for PCM1, PCM2, PCM3 and PCM5, for which we detected solely plasma membrane localization, which is in line with the expectations based on the presence of the CYSTM domain (Venancio and Aravind, 2010) and early reports on PCC1/PCM4 (Mir and Leon, 2014). At this point we cannot exclude that the nucleo-cytoplasmic localization of PCM6-PCM8 detected by us and others (Xu et al., 2018) is due to degradation of the PCM-YFP fusion protein in this experimental setup. Alternative experimental approaches such as biochemical analyses will thus be required to corroborate the subcellular localization of these proteins.

### The function of CYSTM domain-containing PCMs in plant defense

The *PCM4* gene is also known as *PCC1* and has previously been identified as an early-activated gene upon infection with the bacterial pathogen *Pto* carrying the avirulence gene AvrRpt2 and to be controlled by the circadian clock (Sauerbrunn and Schlaich, 2004). Microarray analysis of *npr1*-*1* plants revealed that in addition to several *PR* genes, the expression of *PCC1/PCM4* and *PCM6* were affected in this mutant (Wang et al., 2006). Later, *PCC1/PCM4* was identified to be induced by UV-C light in an SA-dependent manner, potentially playing a role as activator of stress-stimulated flowering in Arabidopsis (Segarra et al., 2010). Transgenic plants carrying the *β-glucuronidase* (*GUS*) reporter gene showed that the expression of *PCC1/PCM4* in the seedling stage was confined to the root vasculature and the stomatal guard cells of cotyledons, but spread to the petioles and the whole limb of fully expanded leaves (Mir et al., 2013). *PCC1/PCM4*-OX lines showed enhanced resistance to *Hpa* (Sauerbrunn 2004; Figure 5A), while RNAi plants were more susceptible to the hemi-biotrophic oomycete pathogen *Phytophthora brassicae* and more resistant to the necrotrophic fungal pathogen *Botrytis cinerea* when compared with wild-type plants (Mir et al., 2013). We confirmed that *PCC1*/*PCM4* overexpression lines are resistant to *Hpa* and extend this finding to additional PCMs: Overexpression of *PCM1, PCM2, PCM3, PCM5, PCM7*, and *PCM8* also provided protection against *Hpa* infection (Figure 5A). This points to a common underlying defense mechanism that is activated by the PCMs, which might be related to an altered membrane environment as we discussed in the previous paragraphs. This mechanism may also be responsible for the enhanced protection against *Pto* infection that we observed by overexpressing *PCM1, PCM2*, and *PCM3* (Figure 5B). The lack of effect on *Pto* of the other *PCM*-OX lines however also points to divergent effects of the different *PCMs*, which is corroborated by the partly distinct DEG sets of the *PCM1*-, *PCM5*-, and *PCM7*-OX lines (Figure 7B). We also assayed the *PCM1, PCM5* and *PCM7* overexpressors for resistance to the biotrophic powdery mildew fungus *Golovinomyces orontii*, but found that these lines displayed the same level of disease development (haustorium formation and macroscopic symptoms) as the wild type, whereas the positive control triple mutant *mlo2 mlo6 mlo12* was highly resistant (Supplemental Figure S3). It may be that the protection mechanism provided by the *PCMs* is not effective against this pathogen species, but it may also be that the species was so virulent that it could have overcome any quantitative resistance accomplished by *PCM* overexpression. Notably, *PCM*-OX lines did not show any morphological abnormalities such as dwarfism, and the RNA-seq data of the *PCM1*-, *PCM5*-, and *PCM7*-OX lines did not reveal any evidence for the constitutive expression of typical defense-related genes (such as *PR* genes) that would explain the enhanced disease resistance of these plant. In the future, it will be of interest to elucidate the yet unrecognized mechanisms that contribute to this phenotype. Conditioned by the antagonistic interplay of defense-associated phytohormones (Leon-Reyes et al., 2010), plants with enhanced resistance to biotrophic pathogens often show enhanced susceptibility to necrotrophic pathogens. For instance infection with hemi-biotrophic *Pseudomonas syringae*, which induces SA-mediated defense, rendered plants more susceptible to the necrotrophic pathogen *Alternaria brassicicola* by suppression of the JA signaling pathway (Spoel et al., 2007). It will thus be also interesting to explore how the *PCM*-OX lines perform upon challenge with necrotrophic pathogens.

### Interplay between immunity and photomorphogenesis

Our transcriptome data revealed that the *PCM1*-, *PCM5*-, and *PCM7*-OX lines were primarily enriched for genes and biological functions related to circadian rhythm, light signaling, and growth and development (Figure 7). The *PCM4*/*PCC1* gene had previously been reported to respond to circadian rhythm and UV-C light, and to have an effect on stress-induced flowering (Segarra et al., 2010). Here, we show that the *PCM1*-, *PCM5*-, and *PCM7*-OX lines exhibit elongated hypocotyl growth compared to wild-type plants (Figure 8). This phenotype is shared with the *hy5 hyh* double mutant, suggesting that the *PCMs* promote photomorphogenesis. Several studies have addressed the connection between plant defense and light signaling; e.g. UV-C induces SA-dependent defenses, and high levels of far red light (as in shade) repress defense responses to both pathogen and insects, as reviewed by Ballaré (2014). A recent paper by Nozue et al. (2018) reported that SA pathway genes are key components of shade avoidance, that *PCM4* and *PCM5* are downregulated by high far red levels, and that these genes have an altered expression level in shade avoidance syndrome mutants. Therefore, a double role for the PCMs in defense and photomorphogenesis is not unexpected. How the PCMs accomplish this dual function is not clear yet. Like discussed earlier, the PCMs might influence membrane structure and activity of proteins that reside in the membrane or that bind to PCMs, like the subunit 5 of the COP9 signalosome (Mir et al., 2013; Mir and Leon, 2014). These diverse effects may independently influence defense and photomorphogenesis, but an interdependence between the two biological processes, where one is a consequence of the other, is also a possibility.

In conclusion, our approach led to the identification of the family of PCM proteins that carry the distinctive CYSTM module, and which have a broad biological impact on plant performance, as shown by the enhanced protection against biotrophic pathogens and the enhanced hypocotyl growth in *PCM*-OX lines. We elucidated some molecular effects of the PCMs by showing that the majority of the PCM members localize to the plasma membrane, that the *PCM* genes are responsive to SA and pathogen challenge, and that overexpression of *PCMs* leads to the induction of genes associated with light responses and development, but not to typical defense-associated responses.

## MATERIAL AND METHODS

### Plant material and cultivation conditions

*Arabidopsis thaliana* wild type accession Col-0, mutant *eds1*-*2* (Bartsch et al., 2006), triple mutant *mlo2-5 mlo6*-2 *mlo12*-*1* (Consonni et al., 2006), *hy5 hyh* (Van Gelderen et al., 2018) and *PCM* overexpression lines were used in this study. For whole plant assays with pathogen infection and SA treatment, the seeds were stratified for 48 h in 0.1% agar at 4^°^C prior to sowing them on river sand that was saturated with half-strength Hoagland nutrient solution containing 10 mM Sequestreen (CIBA-GEIGY GmbH, Frankfurt, Germany). After 2 weeks, the seedling were transferred to 60-mL pots containing a soil:river sand mixture (12:5 vol/vol) that had been autoclaved twice for 1 h. Plants were cultivated in standardized conditions under a 10-h day (75 μmol m^−2^ s^−1^) and 14-h night cycle at 21°C and 70% relative humidity. Plants were watered every other day and received modified half-strength Hoagland nutrient solution containing 10 mM Sequestreen (CIBA-GEIGY GmbH, Frankfurt, Germany) once a week. To minimize within-chamber variation, all the trays, each containing a mixture of plant genotype or treatments, were randomized throughout the growth chamber once a week. For the hypocotyl elongation assay seeds were surface-sterilized and sown on MS plates (8 g l^− 1^ agar and 1 g L^−1^ Murashige and Skoog (Duchefa Biochemie B.V., Haarlem, The Netherlands)). The seeds were stratified in the dark at 4°C for 2-3 days before being moved to a climate chamber with long-day conditions (16 h light : 8 h dark). After 7 days the plates were photographed and hypocotyl length was measured using ImageJ as described previously (De Wit et al., 2016).

The Arabidopsis *PCM* overexpression lines were generated by amplifying the coding sequence of genes *At2g32190* (*PCM1*), *At2g32200* (*PCM2*), *At2g32210* (*PCM3*), *At3g22231* (*PCM4/PCC1*), *At3g22235* (*PCM5*), *At3g22240* (*PCM6*), *At1g05340* (*PCM7*) and *At1g56060* (*PCM8/ ATCYSTM3*) from accession Col-0. The *PCM* genes were part of a recent paper by Xu et al. (2018) and were named differently in the present study, as clarified in Supplemental Table S1. The primers used for cloning are also listed in Supplemental Table S1. The DNA sequence of the PCR fragments was verified and then cloned using Gateway® cloning (Invitrogen) in the pENTR vector, and subsequently in the pFAST-GO2 Gateway® (Shimada et al., 2010) compatible binary vector under control of the CaMV 35S promoter, followed by sequence verification. Binary vectors were transformed into *Agrabacterium tumefaciens* strain C58C1 containing pGV2260, which was used to transform accession Col-0 using the floral dip method (Clough and Bent, 1998). Transformants were selected by growth on ½ MS plates containing DL-Phosphinothricin BASTA, and resistant T_1_ seedlings were transplanted to soil for seed production. T_2_ lines were selected for single insertion of the transgenes using BASTA resistance. Finally, T_3_ seeds were screened for homozygosity using GFP signal in dry seed coating marker. Experiments were performed using homozygous T_3_ or T_4_ seeds.

### RNA-seq library preparation and sequencing

The experimental design of the RNA-seq time series experiment with SA-treated Arabidopsis leaves has been described previously (Hickman et al., 2019). In brief, the rosettes of 5-week-old Arabidopsis accession Col-0 plants were dipped into a solution containing 1 mM SA (Mallinckrodt Baker) and 0.015% (v/v) Silwet L77 (Van Meeuwen Chemicals BV). For mock treatments, plants were dipped into a solution containing 0.015% (v/v) Silwet L77. The sixth leaf (counted from the oldest to the youngest) was harvested from four individual SA-or mock-treated plants at each of the following time points post-treatment: 15 min, 30 min and 1, 1.5, 2, 3, 4, 5, 6, 7, 8, 10, 12 and 16 h. Total RNA was extracted using the RNeasy Mini Kit (Qiagen), including a DNase treatment step in accordance with the manufacturer’s instructions. RNA-seq library preparation and sequencing was performed by UCLA Neuroscience Genomics Core (Los Angeles, CA, USA). Sequencing libraries were prepared using the Illumina TruSeq RNA Sample Prep Kit, and sequenced on the Illumina HiSeq 2000 platform with single read lengths of 50 bases.

For the comparison of the *PCM1-*OX, *PCM5-*OX and *PCM7-*OX lines with wild-type Col-0, two mature leaves (developmental leaf number six and seven) were harvested from two 5-week-old plants per genotype, resulting in two biological replicates. RNA-seq library preparation and sequencing was performed by the Utrecht Sequencing Facility (Utrecht, Netherlands). Sequencing libraries were prepared using the Illumina Truseq mRNA Stranded Sample Prep Kit, and sequenced on the Illumina NextSeq 500 platform with read lengths of 75 bases.

### RNA-seq analysis

Quantification of gene expression from RNA-seq data was performed as described previously (Caarls et al., 2017; Hickman et al., 2017). Reads were mapped to the Arabidopsis genome (TAIR version 10) using TopHat version 2.0.4 (Trapnell et al., 2009) and aligned reads summarized over annotated gene models using HTseq-count (Anders et al., 2015). Genes that were significantly altered over time in response to SA in comparison to the mock treatment were identified using a generalized linear model implemented with the R statistical environment (www.r-project.org). Genes that were differentially expressed between Col-0 and *PCM1-*OX, *PCM5-*OX, or *PCM7-*OX were identified using DESeq2 (Anders and H., 2010; Love et al., 2014).

### Identification of uncharacterized gene families

Protein sequences of the 630 SA-responsive DEGs with unknown/uncharacterized function (based on gene annotations retrieved from TAIR version 10 (retrieved in 2016) were run through OrthoMCL with default parameters (www.orthomcl.org) (Li et al., 2003). JackHMMER (www.ebi.ac.uk/Tools/hmmer/search/jackhmmer) was then used to identify additional paralogs belonging to the groups identified with OrthoMCL. The phylogentic tree of PCM homologs was generated using PLAZA v4.0 (https://bioinformatics.psb.ugent.be/plaza/) with the *PCM1* gene as a query (Van Bel et al., 2018).

### Determination of transcription factor binding motifs

Transcription factor-gene interactions were inferred from DAP-seq (DNA affinity purification sequencing) experiments, which provide the genome-wide binding profiles of in-vitro-expressed TFs (O’Malley et al., 2016). DAP-seq peaks for 349 Arabidopsis transcription factors with a FRiP (fraction of reads in peaks) score ≥5% were retrieved from the Plant Cistrome DB (O’Malley et al., 2016). DAP-seq peaks were used to infer representation of DNA-binding motifs in the promoters of the *PCM* genes. Motifs are grouped according to cognate transcription factor family.

### Coexpression network analysis

The *PCM* coexpression network was obtained using the ATTED-II Network Drawer tool with the Ath-r platform (http://atted.jp/cgi-bin/NetworkDrawer.cgi) (Obayashi et al., 2017) using the *PCM* genes as query genes. Coexpression networks were visualized using Cytoscape v.3.5.1 (Shannon et al., 2003).

### Functional enrichment analysis

GO-term enrichment analysis on gene lists was performed using the GO term finder tool (Boyle et al., 2004). Where indicated, generic GO terms were removed from the analysis by limiting the maximum size of functional categories to 1500 genes.

### Construction of YFP-tagged PCMs and visualization by confocal microscopy

For *in planta* localization experiments, cDNA extracted from Arabidopsis was used to amplify the CDSs without the stop codon of *PCMs* using the primers listed in the Supplemental Table 1. The PCR products containing *att*B sequence were cloned into the Gateway pDONR221 vector, then the resulting entry vectors containing *PCM* genes were recombined into the Gateway expression vector pB7WGY2, which contains the coding sequence of the Venus fluorescent protein (a YFP variant).

Competent cells of *A*. *tumefaciens* were transformed with the Gateway expression vector described in the previous paragraph made for protein localization. Transformed colonies were selected using the antibiotic resistance of vector and with rifampicin carried by *A*. *tumefaciens*. Single colonies were grown for 2 days at 28^°^C in 20-mL LB medium under shaking conditions. After, the OD_600_ was measured, the cells were pelleted and resuspended to a final OD_600_ of 0.5 with a ½ MS medium (Duchefa Biochemie) supplemented with 10 mM MES hydrate (Sigma-Aldrich), 20 g L^−1^ sucrose (Sigma-Aldrich), 200 µM acetosyringone (Sigma-Aldrich) at pH 5.6 and incubated in darkness for at least 1 h. The solutions were used to agroinfiltrated the abaxial side of 4-5-week-old *N*. *benthamiana* leaves using a 1-mL syringe. The plants were left to grow in normal light conditions and after 2 days leaf sections were taken from agroinfiltrated regions and visualized with confocal microscope.

Microscopy was performed using a Zeiss LM 700 (Zeiss, Germany) confocal laser-scanning microscope. Fresh leaf material was prepared on glass slide with cover slip. Excitation of YFP and RFP (plasma membrane FM4-46 dye (Sigma-Aldrich) plus autofluorescence of chlorophyll were done at 488 nm. Light emission of YFP was detected at 493-550 nm and the red signal for the FM4-46 dye at 644-800 nm. Analyses of the images were performed with ZEN lite (blue edition).

### Pathogen cultivation and bioassays

*Hyaloperonospora arabidopsidis* isolate Noco2 (*Hpa* Noco2) spores were harvested from infected (*eds1-*2 mutant) plants, eluted through Miracloth, and diluted in water to 50 spores µL^−1^. For the disease bioassay, 5-week-old plants were spray-inoculated with this spore suspension. Plants were subsequently placed at 100% RH, under short day conditions (9 h light/15 h dark) at 16°C. After 9 days the spores from eight individual rosette plants were harvested in 5-mL of water and the number of spores per milligram of plant tissue (fresh weight of aerial parts) was counted using a light microscope. Spore counts in the mutant and overexpression lines were compared using ANOVA followed by Tukey’s multiple comparison tests.

*Pseudomonas syringae* pv. *tomato* (*Pto*) DC3000 was cultured in King’s B medium supplemented with 50 mg L^−1^ rifampicine at 28°C overnight. Bacteria were collected by centrifugation for 10 min at 4000 rpm, and re-suspended in 10 mM MgSO_4_. The suspension was adjusted to OD_600_=0.0005 and pressure infiltrated into 3 mature leaves of 5-week-old plants with a needleless syringe. After 3 days, leaf discs of 5-mm diameter were harvested from two inoculated leaves per plant, representing a single biological replicate. Eight biological replicates were harvested for each genotype. Subsequently, 500 µL of 10 mM MgSO_4_ was added to the leaf discs, after which they were ground thoroughly with metal beads using a TissueLyser (Qiagen). Serial ten-fold dilutions were made in 10 mM MgSO_4_, and 30 µl aliquots plated onto KB agar plates containing 50 mg mL^−1^ rifampicine. After 48 h of incubation at 28°C, bacterial colonies were counted. Statistical analyses were performed using ANOVA followed by Tukey’s multiple comparison test for means of log_10_-transformed colony counts.

For powdery mildew assays, Arabidopsis plants were inoculated with powdery mildew (*Golovinomyces orontii*) at roughly 2.5 cm rosette size (radius) at four to five weeks after germination. *G*. *orontii* is adapted to infection of Arabidopsis (Kuhn et al., 2016) and was cultivated on susceptible *eds1*-2 plants. Inoculation was conducted by leaf-to-leaf transfer of conidiospores. Leaves from five individual plants were collected at 48 hours post inoculation and bleached in 80% ethanol at room temperature for at least two to three days. Prior to microscopic analysis, fungal structures were stained by submerging the leaves in Coomassie staining solution (100% v/v ethanol acid, 0.6% w/v Coomassie blue R-250; Carl Roth, Karlsruhe, Germany) twice for 15-30 s and shortly washed in tap water thereafter. The samples were analyzed with an Axiophot microscope (Carl Zeiss AG, Jena, Germany). The fungal penetration rate was determined as the percentage of spores successfully developing secondary hyphae over all spores that attempted penetration, visible by an appressorium (Haustorium index). Macroscopic pictures of *G*. *orontii*-infected plants were taken at 12 days post inoculation with a Coolpix P600 camera (Nikon, Tokyo, Japan). Susceptible Col-0 and the fully resistant *mlo2-*5 *mlo6-*2 *mlo12-*1 triple mutant (Consonni et al., 2006) served as positive and negative control, respectively. Haustorium index in the mutant and overexpression lines were compared using ANOVA followed by Tukey’s multiple comparison tests.

## Supporting information

Supplemental Data Set S1

Supplemental Data Set S2

Supplemental Data Set S3

## Funding

CAPES Foundation, Ministry of Education of Brazil (to M.P.M), Netherlands Organization for Scientific Research through the Dutch Technology Foundation (VIDI 11281 to S.C.M.V.W. and VENI 13682 to R.H.), European Research Council (Grant 269072 to C.M.J.P.).

## Supplemental data

**Supplemental Figure S1.**
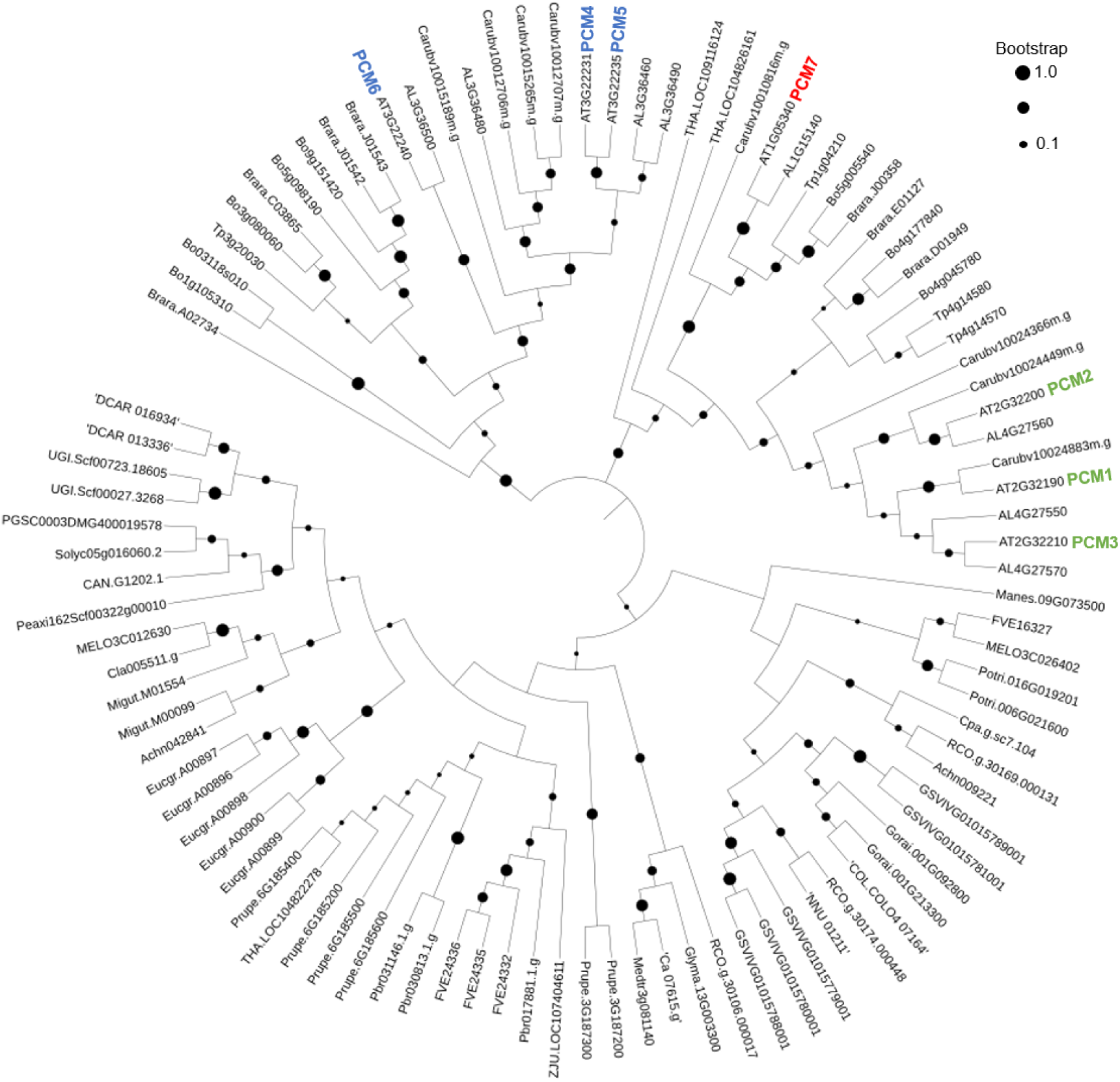
Phylogenetic relationship of closely related homologs of the *PCM* gene family in Arabidopsis and 32 other plant species. The PLAZA platform included the isoform of PCM8 in which the CYSTM domain is excised, for this reason, PCM8 is not included in the phylogenetic tree. Differently sized black dots indicate bootstrap support according to the legend on the top right.

**Supplemental Figure S2.**
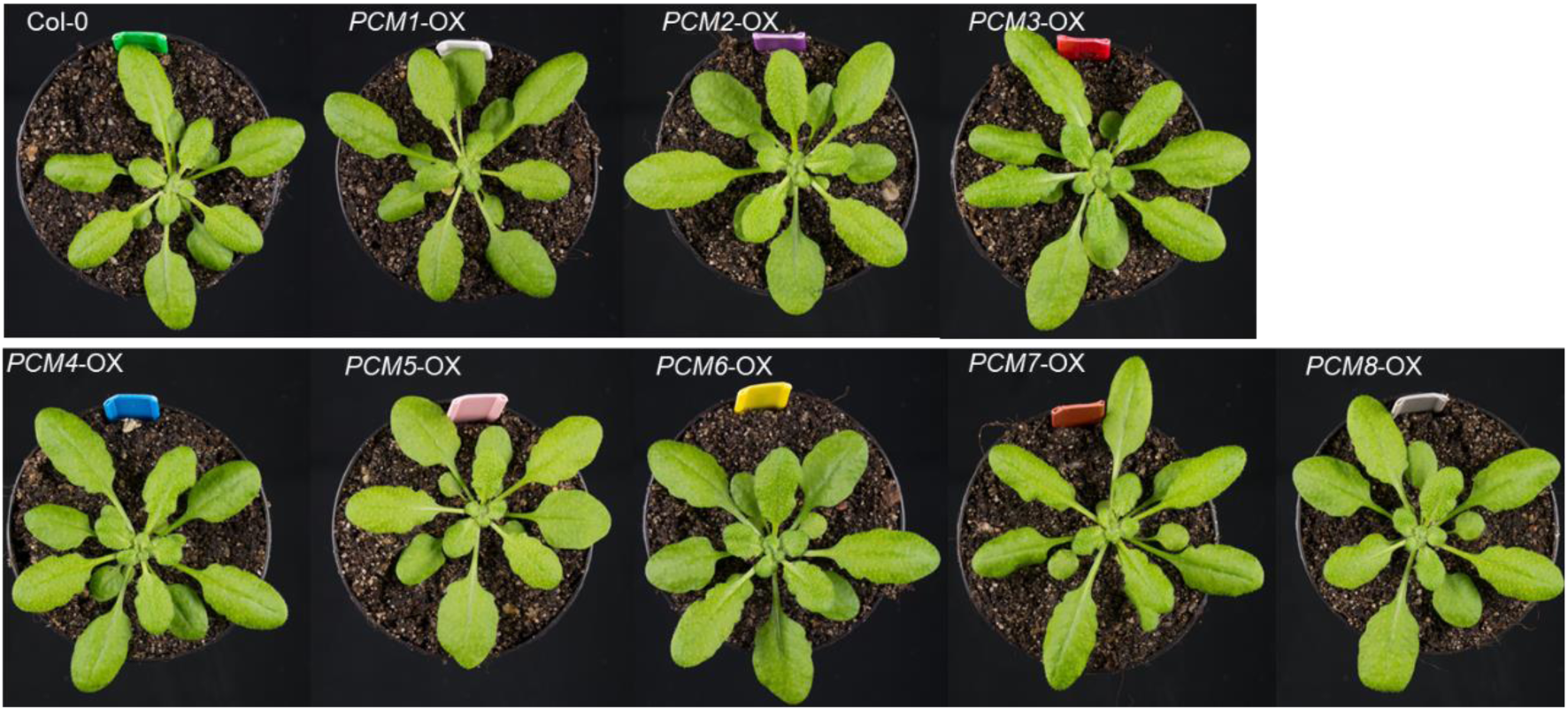
Arabidopsis *PCMs*-OX growth development. Representative photos of 5-week-old plants for each genotype.

**Supplemental Figure S3.**
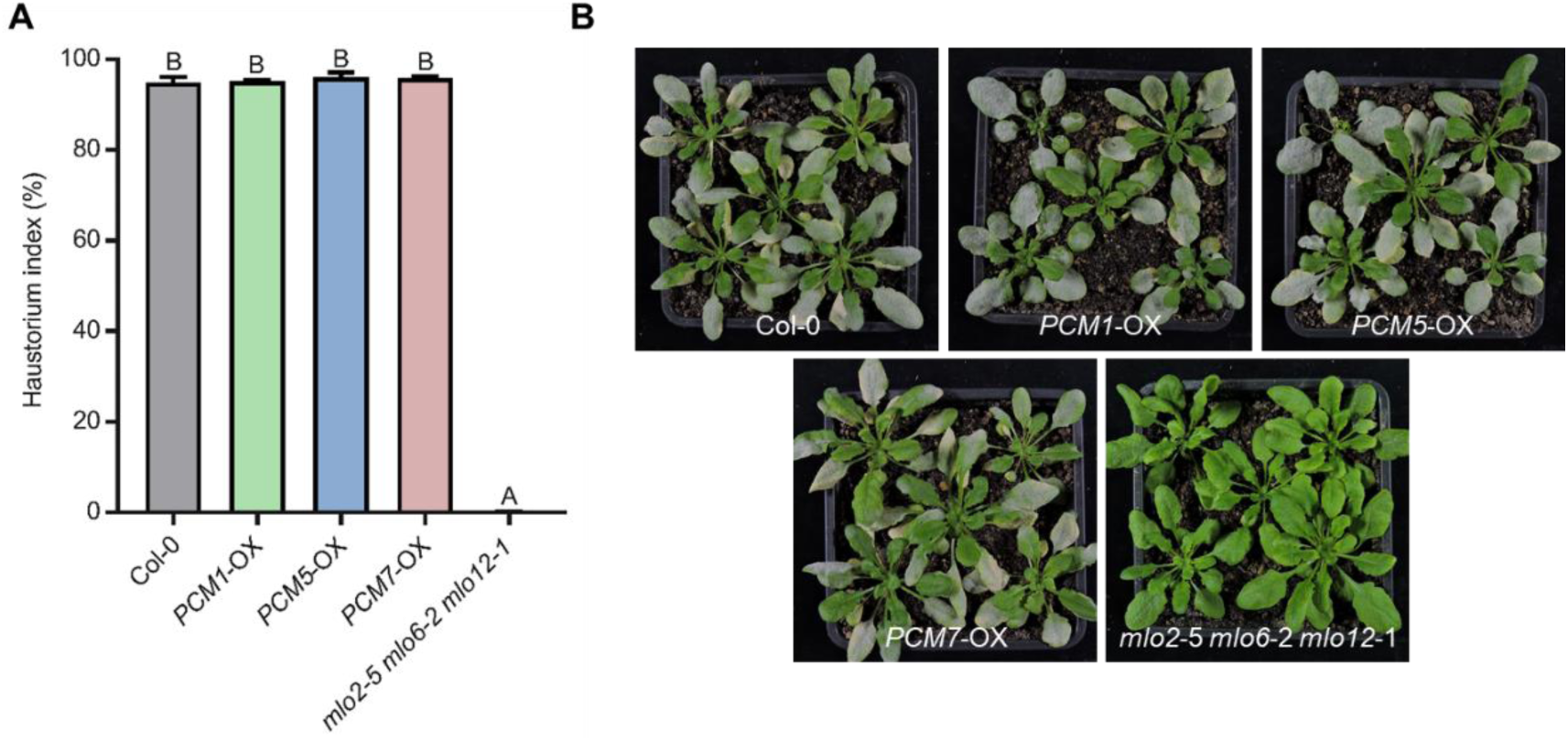
Powdery mildew (*Golovinomyces orontii*) infection phenotypes of *PCM*-OX lines.(**A**), Quantitative analysis of host cell entry (at 48 hours post inoculation) on wild-type Col-0, the fully resistant *mlo2*-5 *mlo6*-2 *mlo12*-1 triple mutant and *PCM1*-OX, *PCM5*-OX and *PCM7*-OX lines. Letters denote significant differences between genotypes (one-way ANOVA, Tukey’s post-hoc test, *P* < 0.05). (**B**), Macroscopic infection phenotypes of the same lines as shown in (**A**) at 12 days post inoculation.

**Supplemental Table S1.**
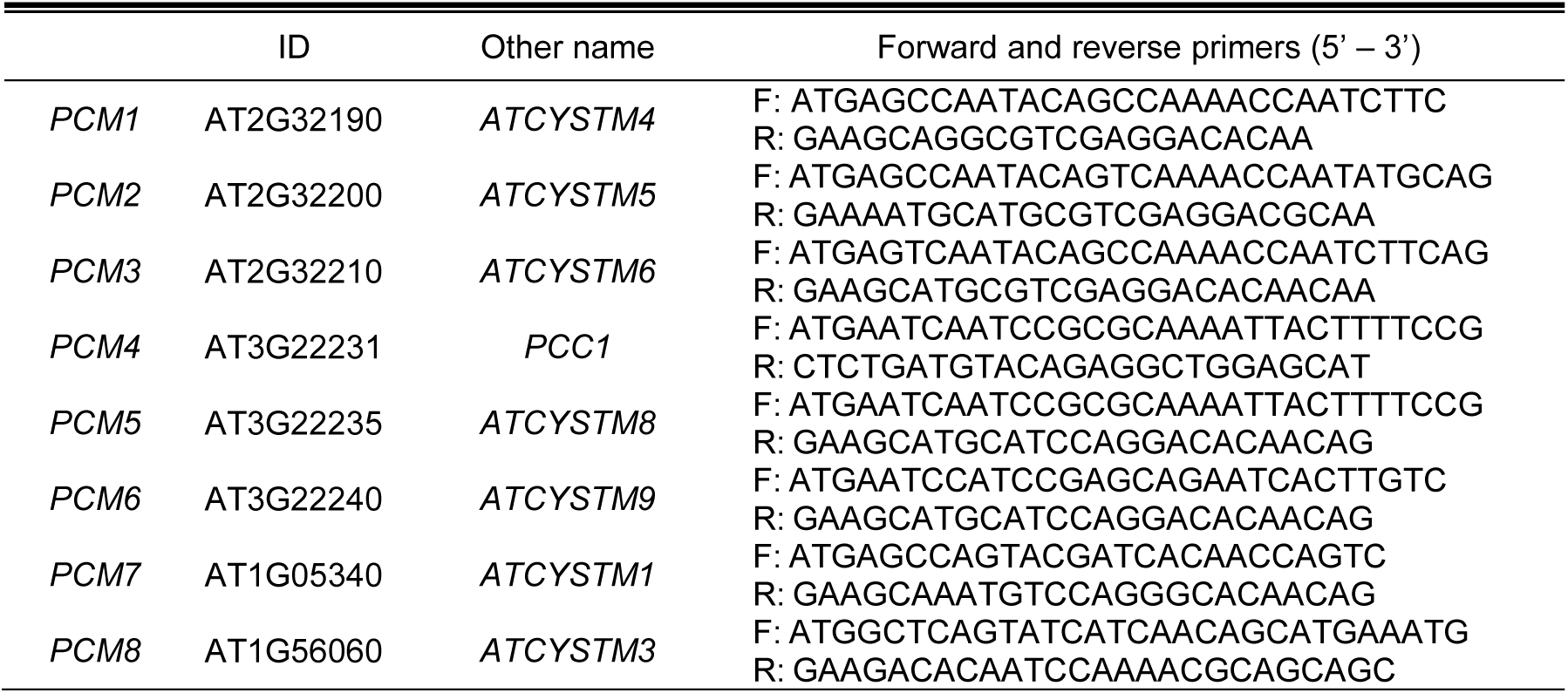
List of *PCM* genes, their AGI numbers (ID) and alternative names. Primer sequences used for cloning.

**Supplemental Data Set S1**. Set of 103 groups of putative homologs among the set of uncharacterized SA-induced genes, identified using OrthoMCL.

**Supplemental Data Set S2**. Genes differentially expressed in *PCM1*-OX, *PCM5*-OX and *PCM7*-OX lines in comparison to wild-type plants.

**Supplemental Data Set S3**. GO terms overrepresented in the DEG sets of *PCM1*-OX, *PCM5*-OX and *PCM7*-OX.

## Notes

### Competing Interest Statement

The authors have declared no competing interest.

